# A Universal Approach to Analyzing Transmission Electron Microscopy with ImageJ

**DOI:** 10.1101/2021.05.27.446036

**Authors:** Jacob Lam, Prasanna Katti, Michelle Biete, Margaret Mungai, Salma AshShareef, Kit Neikirk, Edgar Garza Lopez, Zer Vue, Trace A. Christensen, Heather K. Beasley, Taylor Rodman, Jeffrey L. Salisbury, Brian Glancy, Renata O. Pereira, E. Dale Abel, Antentor Hinton

## Abstract

2

Transmission electron microscopy (TEM) is a scientific research standard for producing nanometer-resolution ultrastructural images of subcellular components within cells and tissues. Mitochondria, endoplasmic reticulum (ER), lysosomes, and autophagosomes are organelles of particular interest to those investigating metabolic disorders. However, there is no clear consensus amongst regarding the best methods for quantifying the features of organelles in TEM images. In this protocol, we propose a standardized approach to accurately measure the morphology of these important subcellular structures using the free program ImageJ, developed by the National Institutes of Health (NIH). Specifically, we detail procedures for obtaining mitochondrial length, width, area, and circularity, in addition to assessing cristae morphology. We further provide methods for measuring interactions between the mitochondria and ER and measuring the length and width of lysosomes and autophagosomes. This standardized method can be used to quantify key features of organelle morphology, allowing investigators to produce accurate and reproducible measurements of organelle structures in their experimental samples.

**SUMMARY:** We discuss a standardized method for measuring and quantifying organelle features using transmission electron microscopy and accessing for interactions between subcellular structures; organelles of focus include mitochondria, endoplasmic reticulum, lysosomes, and autophagosomes.

## 3 INTRODUCTION

Organelles involved in cellular metabolism are some of the most studied subcellular structures. Mitochondria play essential roles in multiple metabolic pathways and are responsible for generating cellular energy. Additionally, mitochondria are involved in apoptosis and cell death pathways, as well as the production and consumption of reactive oxygen species (ROS)^1^. The endoplasmic reticulum (ER) plays a key role in protein and lipid synthesis, as well as in calcium homeostasis^2^. Additionally, autophagosomes and lysosomes work together to degrade and recycle intracellular components and damaged organelles^3^. Organelle dysfunction contributes to the pathogenesis of many diseases, including type II diabetes mellitus, cardiovascular disease, and Alzheimer’s disease^4–6^. Thus, due to their central roles in pathophysiology, studying the structure of these organelles and accurately assessing how they are disrupted under disease conditions is of paramount importance.

Transmission electron microscopy (TEM) is one of the most widely used methods for studying organelle structure. This technique produces nanometer-resolution images by transmitting electrons through an ultrathin section of the sample and magnifying the resulting image using a series of lenses^7^. TEM has enabled researchers to study the ultrastructure of organelles in a wide variety of healthy and diseased cells and tissues, yielding important insights into key cellular processes and disease development^8^.

However, several limitations remain to using TEM when studying organelles. While overall organelle structures can be readily determined from TEM images, quantifying changes in organelle morphology presents a substantial challenge. Further, there has been no clear consensus regarding the most effective methods for measuring organelle features from TEM images. To address this issue, we propose a standardized approach for analyzing TEM images with the image analysis tool ImageJ, which was developed by the National Institutes of Health (NIH). We outline methods for measuring features of important organelles, including mitochondria, ER, lysosomes, and autophagosomes. In addition, we provide strategies for assessing how these organelles interact with one another. By using a standardized method to quantify organelle morphology as proposed by this study, investigators can generate accurate and easily reproducible measurements of organelle features, thereby increasing the precision and relevance of their data.

## 4 PROTOCOL

1. *Installing and Preparing ImageJ Software for Analysis*
  1.1. Download ImageJ software from the NIH website (https://imagej.nih.gov/ij/download.html).
  1.2. Install and open the ImageJ software.
  1.3. Open the Region of Interest (ROI) Manager, which is used to record and keep track of measurements, by selecting **Analyze** ⟶ **Tools** ⟶ **ROI Manager** (Figure 1A).
  1.4. Click on **Analyze** ⟶ **Set Measurements** to input the measurements that ImageJ will make, such as area, circularity, and perimeter (Figure 1A).
  1.5. Drag the image to be analyzed (TIFF or DM3 file) directly into ImageJ (Figure 1B).
    1.5.1. Alternatively, click **File** ⟶ **Open** to open the selected image.
  1.6. Considerations
    1.6.1. For accuracy and reproducibility, ensure that each image contains a scale bar, length, and image magnification. The scale bar and bar length are important for setting the appropriate units within the Image J settings.
    1.6.2. Quantification of samples should be performed by three individuals in a randomized and blinded manner to ensure an unbiased approach.
2. *Analyzing Mitochondrial Morphology*
  2.1. Measuring mitochondrial area, circularity, and perimeter.
    2.1.1. Click on **Freehand Selections** on the toolbar to access the **Freehand** tool.
    2.1.2. Trace the outline of the entire cell. To store the measurement, click **Add** on the ROI Manager. This will be used to normalize later measurements.
    2.1.3. Trace the outer mitochondrial membrane of each mitochondrion. Add the shape to the ROI Manager (Figure 2A).
    2.1.4. Click on **Measure** in the ROI Manager to obtain the measurements for each shape.
  2.2. Measuring mitochondrial length and width
    2.2.1. Click on **Straight, Segmented, or Freehand Lines** on the toolbar and select 4.1 **Straight Line**.
    2.2.2. Draw a straight line along the major and minor axis of each mitochondrion. Add the measurements to the ROI Manager (Figure 2A).
    2.2.3. Click the **Measure** function to obtain lengths and widths.
3. *Analyzing Cristae Morphology*
  3.1. Split each image into four quadrants and randomly pick two quadrants to analyze; use the same quadrants for all images.
  3.2. Using the **Freehand** tool in ImageJ, outline the outer membrane of a mitochondrion. Add the measurement to the ROI Manager.
  3.3. Trace the outline of each crista within the mitochondrion. Add each measurement to the ROI Manager (Figure 2B).
  3.4. Click on **Measure** in the ROI Manager to obtain the area of each crista, and cristae surface area is the sum of the areas of all the cristae in a single mitochondrion.
  3.5. To calculate cristae volume density, cristae surface area is divided by the area of the mitochondrion.
  3.6. To determine the cristae number, count the number of cristae in each mitochondrion. Alternatively, you can use the number of cristae measurements obtained earlier.
  3.7. To assign a mitochondrion a cristae score, evaluate cristae abundance and form. Assign a score between 0 and 4 (0 – no sharply defined crista, 1 – greater than 50% of mitochondrial area without cristae, 2 – greater than 25% of mitochondrial area without cristae, 3 – many cristae but irregular, 4 – many regular cristae)^9^.
  3.8. Considerations
    3.8.1. Average cristae measurements in cells are based on 1000 mitochondria from 100 cells or 10 mitochondria per cell from 2,000 cells. For tissue samples, average measurements are based on three independent animals with 330 mitochondria measured from at least 10 images.
4. *Analyzing Mitochondria-ER Contacts (MERCs)*
  4.1. Split each image into four quadrants and randomly pick two quadrants to analyze; use the same quadrants for all images.
  4.2. Identify a Mitochondrion-ER contact (MERC), a mitochondrion in close contact with the ER membrane, in your TEM image.
  4.3. Using the **Freehand Selections** tool, trace both the outer membrane of the mitochondrion and the ER membrane at a contact point (Figure 2C).
  4.4. To obtain the MERC contact length, click on the **Straight, Segmented, or Freehand Lines** tool on the toolbar, select **Freehand Line**, and draw a line spanning the length of the contact site (Figure 2C). Add the measurement to the ROI Manager and use the **Measure** function to determine the length of the contact.
  4.5. To measure the MERC distance, use the **Freehand Line** tool to draw a line between the two organelles (Figure 2D). Add the measurement to the ROI Manager and use the **Measure** function to determine the distance.
  4.6. To calculate percent coverage, divide the contact length by either the mitochondrial surface area (mitochondrial percent coverage) or ER surface (ER percent coverage), and multiply the value by 100 to obtain a percentage.
  4.7. Similar measurements can be made for other organelle interactions using the same steps detailed here.
  4.8. Considerations
    4.8.1. Average MERC changes in cells should be based on 100 to 400 images. For tissue samples, measurements should be based on three independent animals with 100 to 400 images per animal.
    4.8.2. Average MERC distance and length in cells should be based on 50 to 100 images. For tissue samples, measurements should be based on three independent animals with 10 to 15 images per animal
5. *Analyzing Lysosomes and Autophagosomes*
  5.1. Click on the **Freehand Selections** to access the **Freehand** tool.
  5.2. Trace the outline of the entire cell. Click **Add** on the ROI Manager. This will be used to normalize later measurements.
  5.3. Trace the membrane of each lysosome and autophagosome. Add the shape to the ROI Manager (Figure 2E). Click **Measure** in the ROI Manager to obtain measurements for the area.
  5.4. To obtain length and width, use the **Straight Line** tool to draw a line down the major and minor axes of each organelle. Add the measurements to the ROI Manager and use the **Measure** function to obtain numerical values for each measurement.
6. *Pseudo-coloration of Organelles* (Supplementary Figure 1).
  6.1. Install and open Adobe Photoshop.
  6.2. Click **File** ⟶ **Open** and select the saved ImageJ export file.
  6.3. Under the **Layers** menu click **New** ⟶ **Layer** and ensure the new layer is selected and above the layer with the image.
  6.4. Outline the organelle (Figure 3).
    6.4.1. On the left tool sidebar, select the **Freeform Pen Tool**.
    6.4.2. Use tiny increments to slowly outline.
    6.4.3. If mistakes are made, use the eraser tool to fix them.
    6.4.4. Ensure one continuous line is used for the outline of organelles.
    6.4.5. Right click the line drawn and click **Stroke Path**, set to pencil or paintbrush to create a continuous and clean line (Figure 3A).
    6.4.6. Right click the original line drawn and click **Delete Path** to remove the original line and only be left with the clean version (Figure 3B).
  6.5. Clean up the outline
    6.5.1. On the left tool sidebar, select the **Eraser**.
    6.5.2. Use the eraser to manually erase parts where the lines are uneven or unrepresentative of the organelle.
  6.6. Under the **Layers** menu click **New** ⟶ **Layer** and ensure the new layer is selected and below the layer with the outline.
  6.7. Color the organelle.
    6.7.1. On the left hand side, select the **Paint Brush** tool and at the bottom select a color (Figure 3C).
    6.7.2. Use the mouse to color in the area outlined, this does not need to be precise as it will later be adjusted (Figure 3D).
    6.7.3. To fix uneven coloring, once fully colored, for each organelle select the **Magic Wand**. Go to the outline layer and click onto each organelle to cause the coloration to fit evenly inside the outline (Figure 3E).
    6.7.4. Delete the Color layer, so the only layer left is the Outline layer with the new colors and the background image (Figure 3F).
  6.8. Considerations
    6.8.1. During outlining the organelle, one can do small lines and create a stroke path for each line, this increases accuracy but will take a longer time.
    6.8.2. It is recommended that one saves continuously during this process as it is easy to begin working on the wrong layer.
    6.8.3. During the coloration process, so long as the outline is continuous, one can use the **Magic Wand** from the tool menu to instantly fill in the area with one color.
    6.8.4. Organelles can have colored outlines only if the coloration step is skipped and during the outlining step another color besides black is selected for each organelle (Figure 3).
  6.9. Click **File** ⟶**Save As…** and save as a desired export file for further analysis, if necessary.

**Figure 1.**
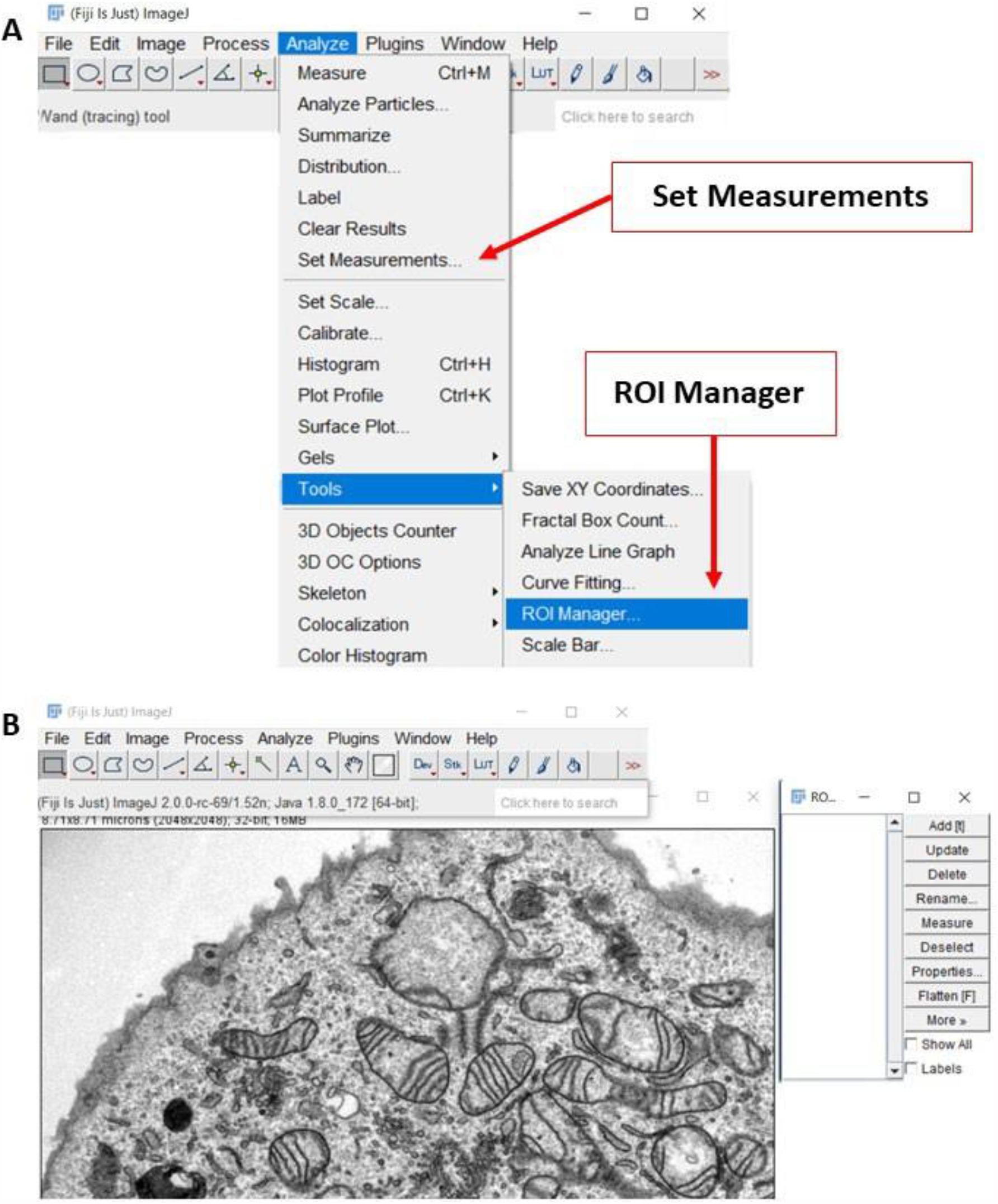
Preparing ImageJ software for image analysis. (**A**) On the ImageJ toolbar, the “Analyze” menu contains many of the settings and tools needed for this analysis method. **(B)** Representative screenshot of a transmission electron microscopy (TEM) image that is ready to be analyzed.

**Figure 2.**
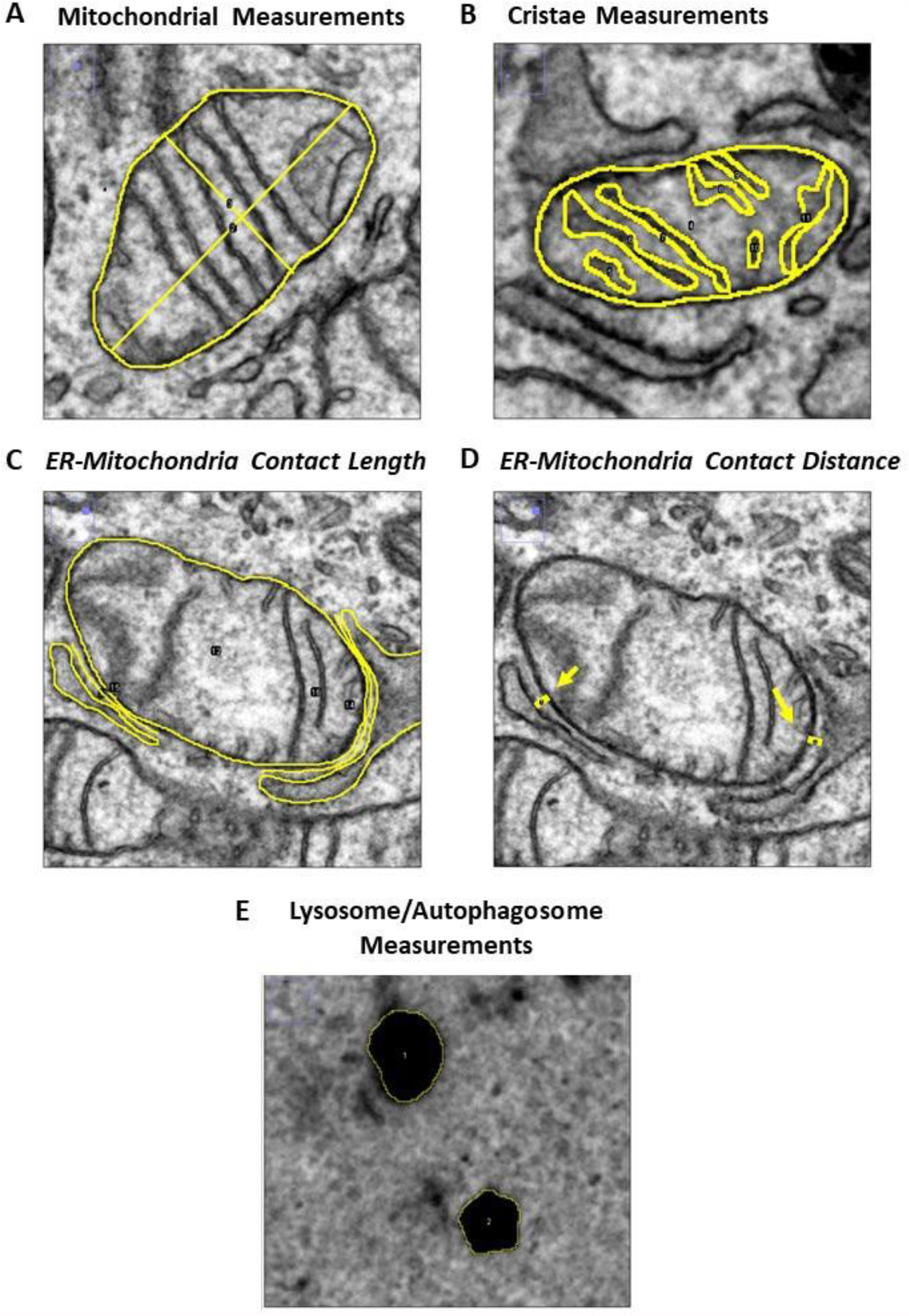
Analyzing organelle morphology with ImageJ. (**A**) Representative TEM image illustrating how to measure mitochondrial length, width, area, and circularity. (**B**) Representative TEM image illustrating how to obtain cristae measurements (yellow). (**C, D**) Representative TEM images illustrating how to determine ER-mitochondria contact length and ER-mitochondria contact distance (yellow). (**E**) Representative TEM image illustrating how to obtain lysosome and autophagosome measurements.

**Figure 3.**
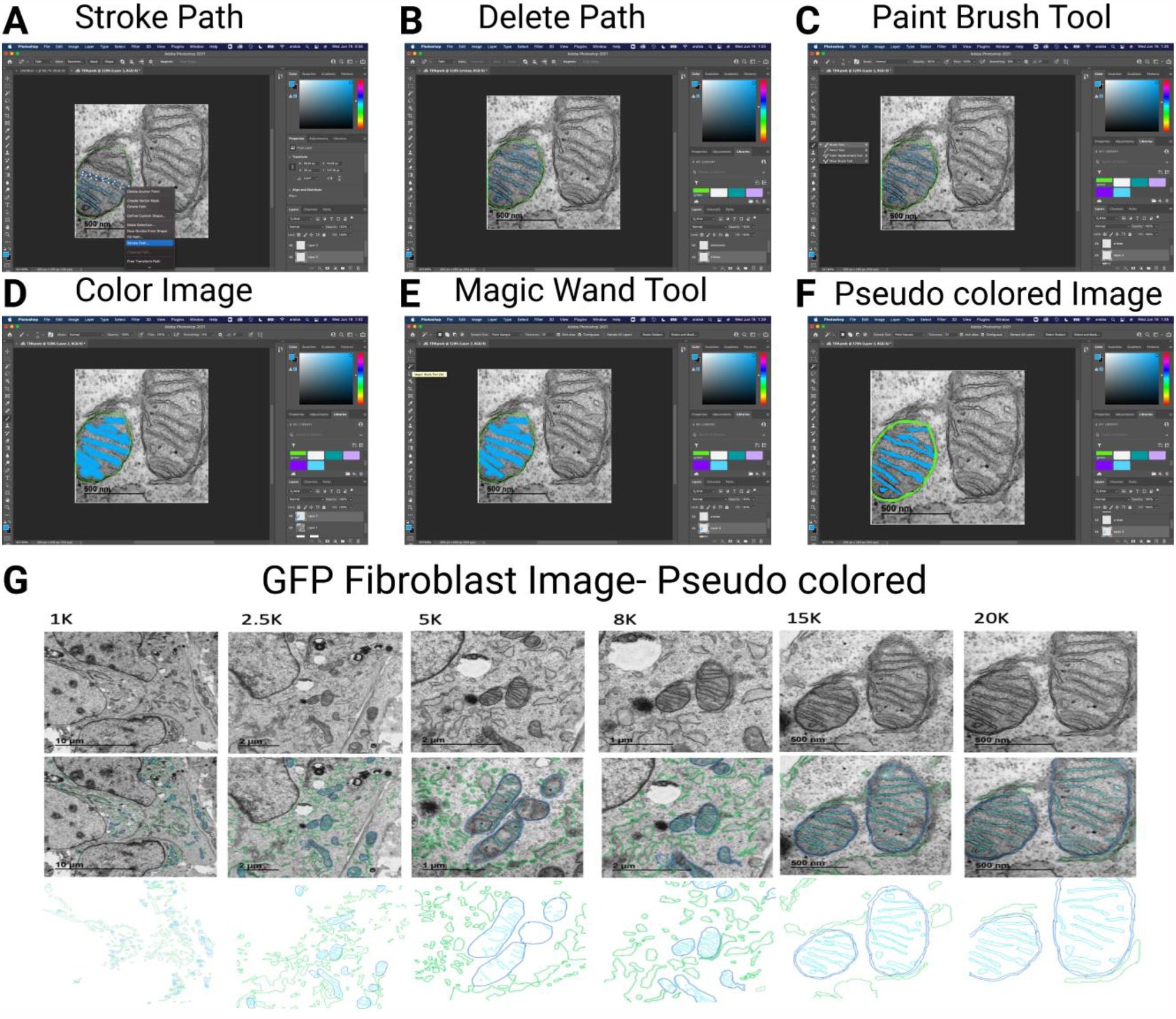
Pseudo-coloration of organelles with Adobe Photoshop. Organelles at various magnifications can be ported from ImageJ to Adobe Photoshop for outlining and coloration. This shows cristae segmentation and coloration with blue and mitochondria with green. Representative images are shown for each magnification to the left. Coloration allows for the clean presentation and easy visual analysis of TEM images. (**A**) Image showing the utilization of the stroke path feature in Adobe for reducing pixelated edges. (**B**) Image showing a mitochondrion that has been outlined in color in Adobe Photoshop. (**C**) Image showing selection of brush tool from the toolbar in Adobe Photoshop. (**D**) Image showing the pre-correction colored in organelle in Adobe Photoshop. The initial coloration does not need to be exact. (**E,F**) The selection and application of the Magic Wand tool fits the colors into the previously created outlines and allows for the pseudo-coloration of organelles. (G) The representative figure at varying magnifications of the mitochondria in fibroblast cells.

## 5 REPRESENTATIVE RESULTS

This protocol standardizes an approach to obtaining accurate and reproducible measurements of organelle morphological features. Below, we discuss findings based on this TEM image analysis approach.

### 5.1 OPA-1 Knockdown Decreases Mitochondrial Area and Disrupts Cristae Structure

Optic atrophy protein-1 (OPA1) is an inner mitochondrial membrane protein that plays an essential role in inner mitochondrial membrane fusion, in concert with mitofusin proteins MFN1 and MFN2 that mediate outer membrane fusion^10^. Previous studies have demonstrated that OPA1 knockdown inhibits mitochondrial fusion and increases the number of fragmented mitochondria^11^. Using our proposed method outlined above, we confirmed this finding in mouse skeletal muscle tissue (Figure 4A). Specifically, we showed that knocking down *Opa1* significantly decreases the mean mitochondrial area for the peri-nuclear mitochondria cluster (Figure 4B). Additionally, we observed an increase in both the number of mitochondria per square micron and the average circularity index (Figure 4C and 4D). These findings suggest that loss of OPA1 decreases mitochondrial fusion and increases fission events, consistent with previous studies.

**Figure 4.**
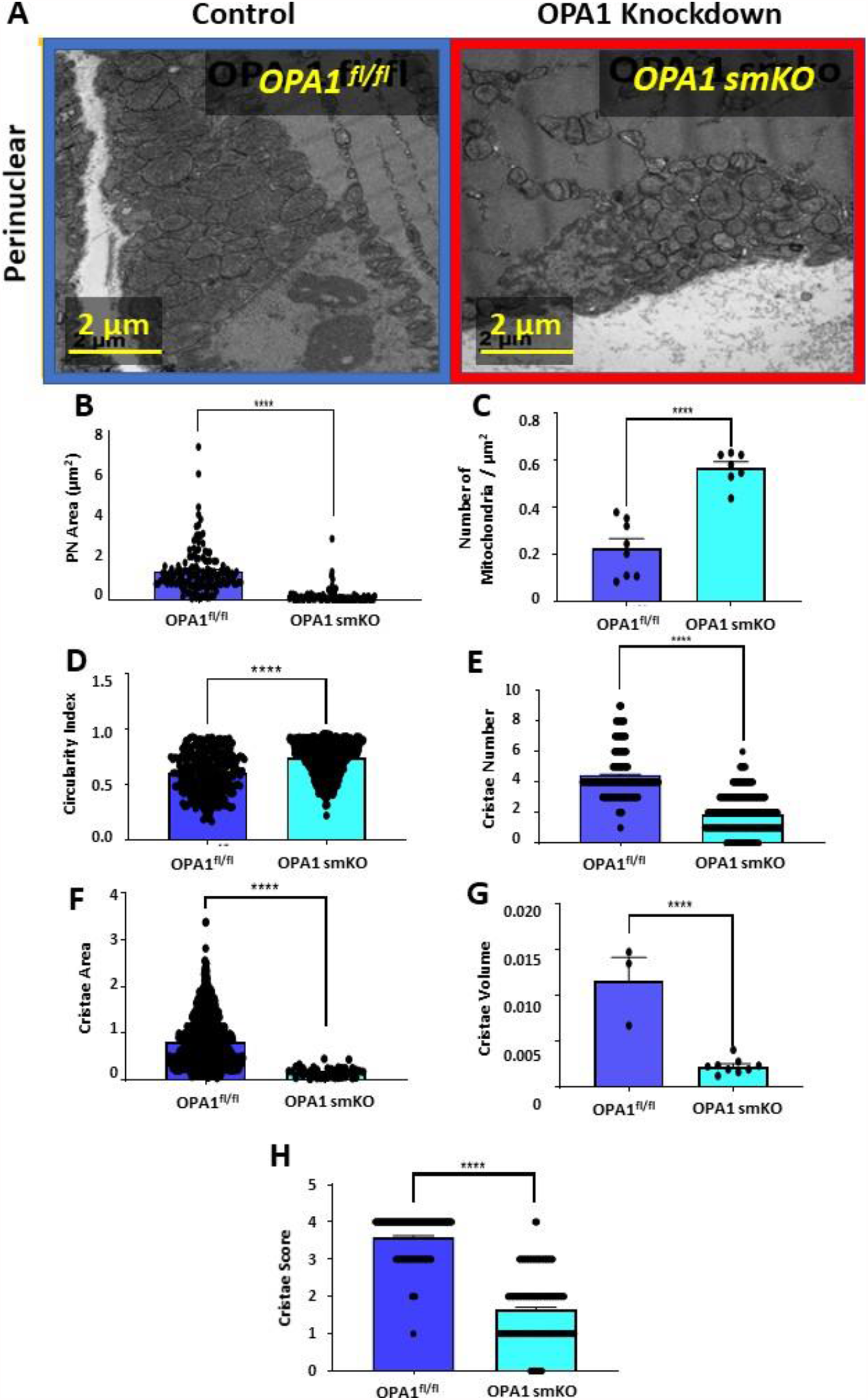
Knockdown of optic atrophy protein-1 (OPA1) decreases mitochondrial area and disrupts cristae structure. (**A**) Representative TEM images of mitochondria in mouse skeletal muscle tissue from OPA1 knockdown (red outline) and control (blue outline) tissue. (**B**) Quantification of perinuclear mitochondrial area, (**C**) number of mitochondria per square micron, and (**D**) mitochondrial circularity index. (**E**) Quantification of cristae number, (**F**) cristae area, (**G**) cristae volume, and (**H**) cristae score. Statistical significance is indicated by asterisks; *, **, ***, **** indicate *p*≤0.05, *p*≤0.01, *p*≤0.001, and *p*≤0.0001, respectively.

Beyond its role in mitochondrial dynamics, OPA1 plays a direct role in cristae remodeling^12^. To assess the effect of OPA1 on cristae morphology, we used our TEM analysis protocol to quantify cristae changes in OPA1-knockdown skeletal muscle tissue. We confirmed disruption of cristae morphology, and reductions in the average number of cristae per mitochondrion, average cristae surface area, and cristae volume density in OPA1-knockdown cells (Figures 4E – 4G). Furthermore, the cristae score was reduced in OPA1-knockdown cells relative to the control group (Figure 4H). These results confirm that OPA1 is essential for cristae remodeling and that loss of OPA1 disrupts normal cristae structure.

### 5.2 Thapsigargin Treatment Increases MERCs

The interaction points between the ER and mitochondria, commonly referred to as mitochondria-ER contacts (MERCs), are increasingly becoming a focus of study in cellular biology. In particular, emerging research has implicated defective MERCs in multiple age-associated diseases, such as neurodegenerative diseases, metabolic syndrome, and cardiovascular diseases^13^. To test our method of quantifying MERCs, we measured the effect of thapsigargin treatment on MERCs in murine cells (Figure 5A). Thapsigargin is a sarco-ER Ca^2+^-ATPase (SERCA) inhibitor that induces ER stress^14^. ER stress inducers like thapsigargin cause mitochondria to move closer to the ER^15^. In our analysis, we found that thapsigargin treatment greatly decreases the distance between mitochondria and the ER compared with the control cells (Figure 5B). The average contact length is also significantly increased in treated cells (Figure 5C). These findings suggest that thapsigargin treatment influences MERC structure by increasing the degree of contact between mitochondria and the ER and decreasing the distance between the two organelles, as previously reported^15^.

**Figure 5.**
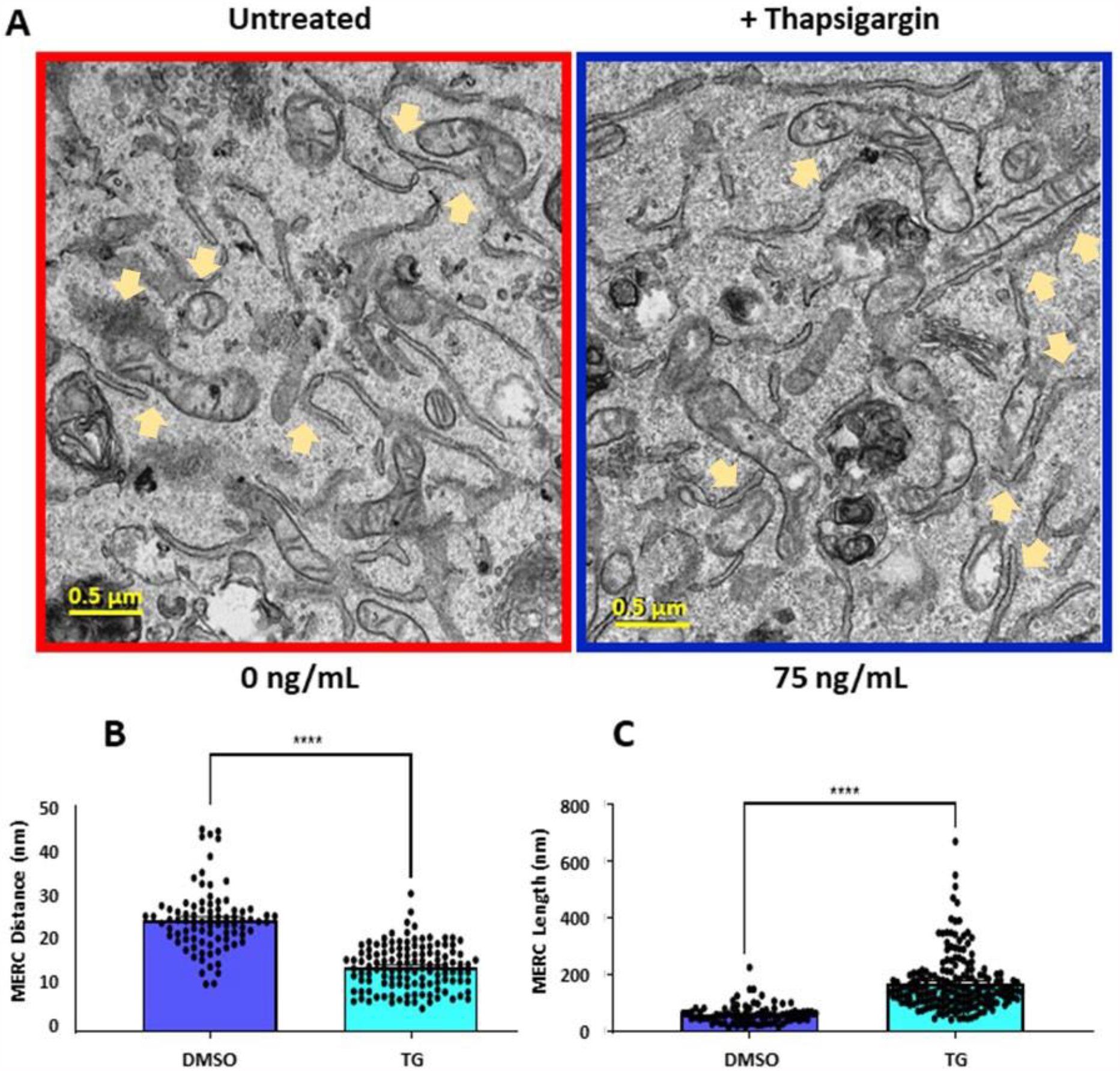
Thapsigargin treatment increases mitochondria-endoplasmic reticulum contacts (MERCs). (**A**)Representative TEM images of mouse skeletal muscle from untreated (blue outline) and (red outline) thapsigargin-treated cells. (**B**) Quantification of MERC distance and (**C**) MERC length. Statistical significance is indicated by asterisks; *, **, ***, **** indicate *p* ≤0.05, *p* ≤0.01, *p* ≤0.001, and *p* ≤0.0001, respectively. MERCs appear at orange arrows.

### 5.3 Thapsigargin Treatment Alters Lysosome and Autophagosome Morphology

We also investigated morphological changes in lysosomes and autophagosomes in response to thapsigargin treatment (Figure 6A). Using our TEM image analysis protocol, we found that both mean lysosomal area and the number of lysosomes per square micron are significantly increased in response to thapsigargin treatment (Figure 6B and 6C). Mean autophagosome area and the number of autophagosomes per square micron are also significantly increased in thapsigargin-treated cells (Figure 6D and 6E). Thapsigargin inhibits ER function and promotes cellular stress. These findings support a model in which cell-stress induced organellar damage induces the activity of the recycling machinery, including lysosomes and autophagosomes, to break down damaged organelles. Thus, the morphological changes detected and quantified here by TEM are consistent with the expected effects of thapsigargin treatment ^15,16^.

**Figure 6.**
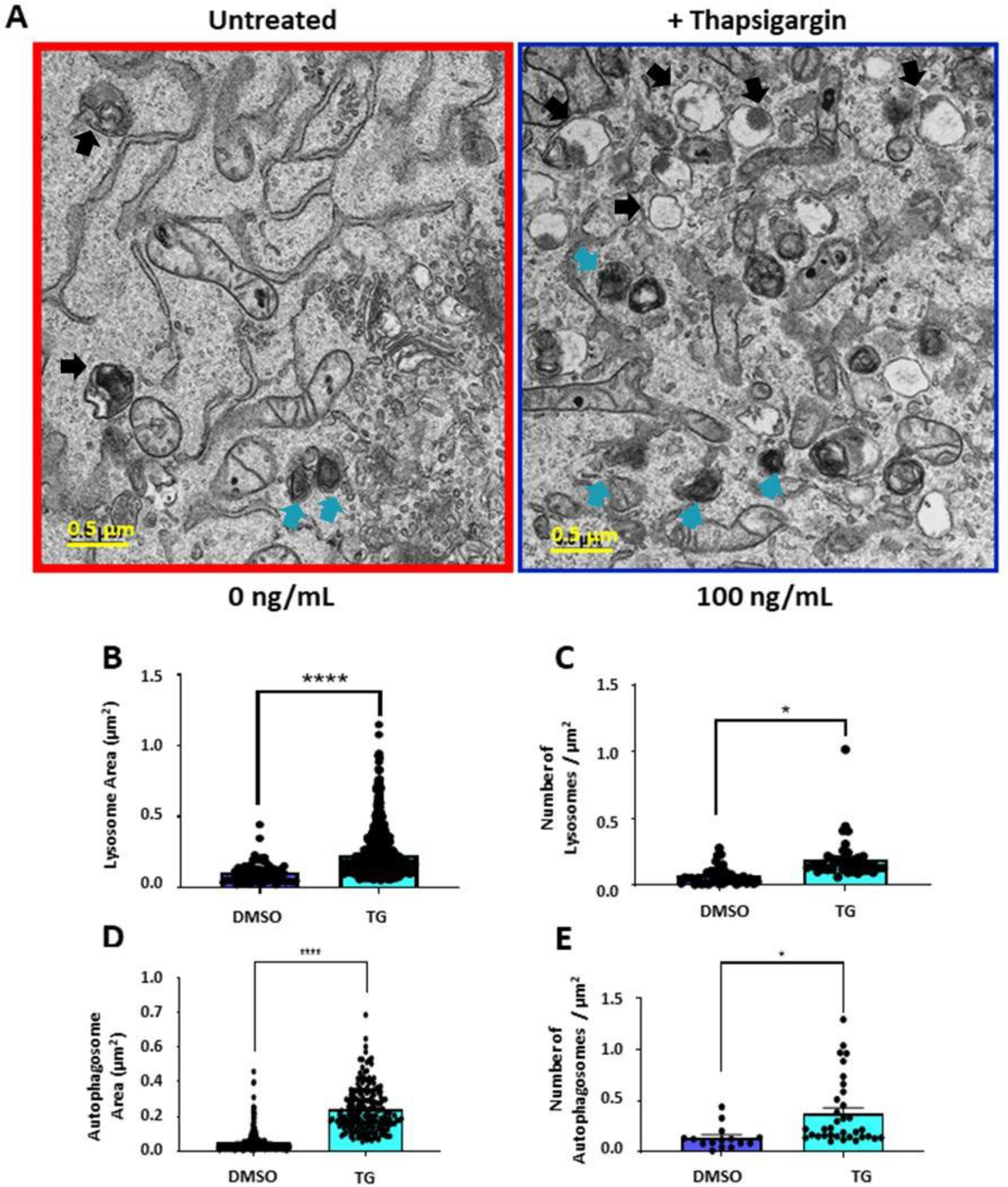
Thapsigargin alters lysosome and autophagosome morphology. (**A**) Representative TEM images of lysosomes and autophagosomes in mouse skeletal muscle from untreated (blue outline) and (red outline) thapsigargin-treated cells. (**B**) Quantification of lysosome area in each treatment group. (**C**) Quantification of the number of lysosomes per square micron. (**D**) Quantification of autophagosome area. (**E**) Quantification of the number of autophagosomes per square micron. Statistical significance is indicated by asterisks; *, **, ***, **** indicate *p* ≤0.05, *p* ≤0.01, *p* ≤0.001, and *p* ≤0.0001, respectively. Autophagosomes appear at black arrows and blue arrows for lysosomes.

## 6 DISCUSSION

TEM is one of the most powerful tools for assessing organelle morphology in cells and tissues. However, to our knowledge, there is no standardized protocol for quantitatively assessing organelle morphology with TEM. Herein, we describe a straightforward approach for accurately quantifying organelle features from TEM images, which can easily be reproduced in all cell and tissue types. Using this method, we verified results from multiple previous studies that assessed how manipulation of mitochondrial and ER proteins using genetic and pharmacological approaches alters organelle morphology. On a broader scale, this method can be applied to any field of study and cell type where organelle structure is a focus. For example, mitochondria play a role in many complex diseases, including type II diabetes, cardiomyopathy, and Alzheimer’s disease^4,6^. Therefore, understanding how these diseases impact mitochondrial structure determined by TEM in conjunction with this precise methodology may lead to a greater understanding of their pathophysiology.

One critical step for successfully implementing this method is to ensure that the image to be analyzed is clear enough to identify specific structures. Several organelles, including the mitochondria, have distinguishing characteristics and are easily observable even at lower magnifications. However, finer structures, such as the cristae within mitochondria, may not be so readily apparent. To address this, higher magnifications of 2,500x or even 5,000x should be used to better resolve organelles of interest.

Although TEM is a useful and powerful tool for assessing organelle structure in cells and tissues, this method of structural analysis has some limitations that may impact our proposed method. Different investigators may possess varying levels of proficiency at using the tools of ImageJ, especially the freehand tools, which could lead to variability in the data or the amount of time required to complete the analysis. However, most investigators should be able to take advantage of this method with practice and the analysis can be completed remotely. Another limitation is that samples must be sliced into ultrathin segments to be properly imaged, meaning three-dimensional characteristics cannot be fully captured by this technique. Thus, features such as organelle volume or organelle interactions in three dimensions cannot be reliably assessed without using alternative methods. In such cases, more advanced techniques, including focused ion beam scanning electron microscopy (FIB-SEM) or serial block-face scanning electron microscopy (SBF-SEM), can be used to image subcellular structures in three dimensions, although accurate quantification is extremely time-consuming^16^. A third limitation of TEM and our analysis method is the lack of live imaging. Due to the requirement of fixing samples in resin, TEM captures only a single snapshot of a sample in time. Consequently, information about dynamic characteristics, such as how organelles move in the cell and change over time cannot be obtained through this approach. Therefore, other techniques, such as fluorescent staining or proximity ligation assay, might be needed to complement this approach^17^. Despite these limitations, our TEM analysis method represents a powerful tool for accurately assessing and quantifying organelle morphology, with the potential for broad application in the study of metabolic disorders and other diseases associated with organelle disruption.

## 7. ACKNOWLEDGEMENTS

We also would like to thank our undergraduate colleagues Benjamin Kirk and Innes Hicsasmaz for helping to optimize the analysis technique. This work was supported by NIH grants R01HL108379 and R01DK092065 to E.D.A, as well as the T32 HL007121, UNCF/BMS EE Just Faculty Fund, the Burroughs Welcome Fund CASI Award, and the Ford Foundation to AHJ.

## 8 DISCLOSURES

The authors have no disclosures to report.

---

**Supplementary Figure 1:**
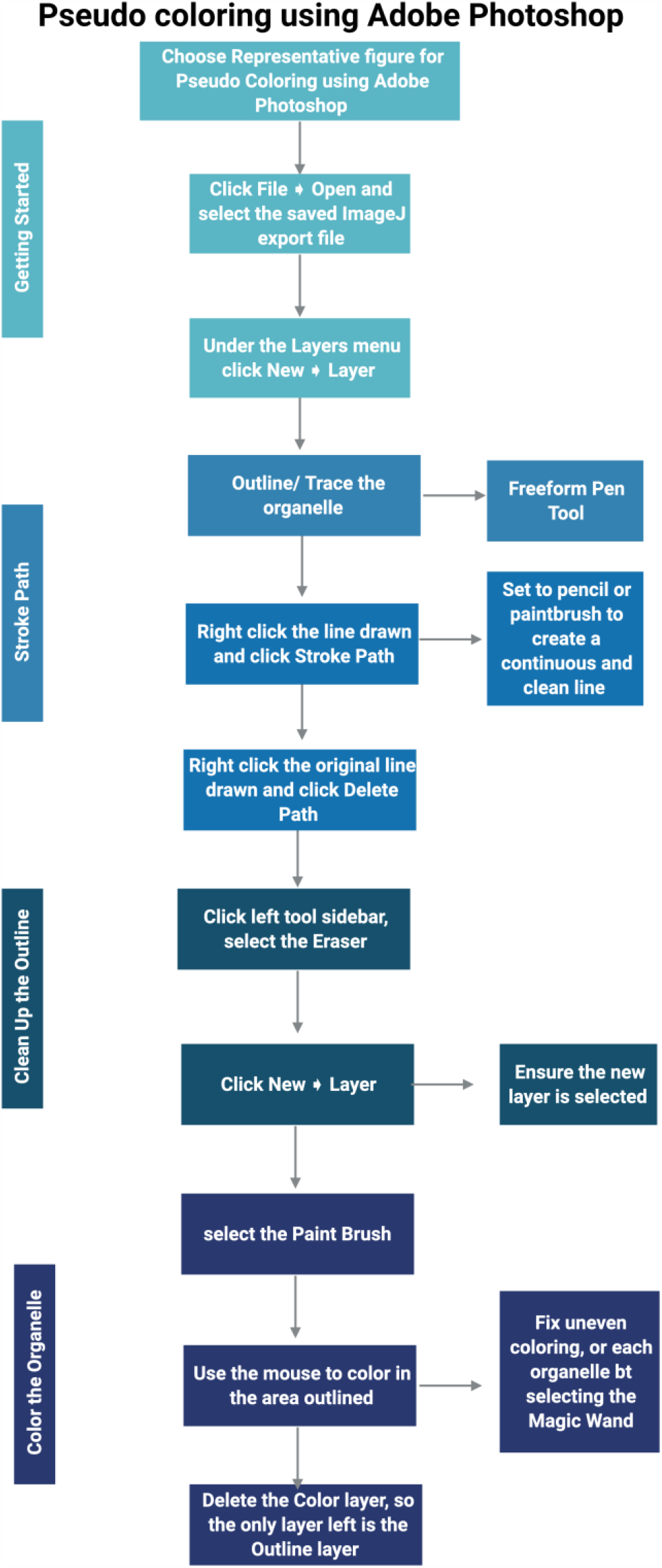
The flowchart representation of Pseudo coloring of TEM images using Adobe Photoshop.

## Notes

### Competing Interest Statement

The authors have declared no competing interest.

